# Protein phosphatase 1 down regulates ZYG-1 levels to limit centriole duplication

**DOI:** 10.1101/093492

**Authors:** Nina Peel, Jyoti Iyer, Anar Naik, Michael P Dougherty, Markus Decker, Kevin F O’Connell

## Abstract

In humans perturbations of centriole number are associated with tumorigenesis and microcephaly, therefore appropriate regulation of centriole duplication is critical. The *C. elegans* homolog of Plk4, ZYG-1, is required for centriole duplication, but our understanding of how ZYG-1 levels are regulated remains incomplete. We have identified the two PP1 orthologs, GSP-1 and GSP-2, and their regulators I-2^szy-2^ and SDS-22 as key regulators of ZYG-1 protein levels. We find that down-regulation of PP1 activity either directly, or by mutation of *szy-2* or *sds-22* can rescue the loss of centriole duplication associated with a *zyg-1* hypomorphic allele. Suppression is achieved through an increase in ZYG-1 levels, and our data indicate that PP1 normally regulates ZYG-1 through a post-translational mechanism. While moderate inhibition of PP1 activity can restore centriole duplication to a *zyg-1* mutant, strong inhibition of PP1 in a wild-type background leads to centriole amplification via the production of more than one daughter centriole. Our results thus define a new pathway that limits the number of daughter centrioles produced each cycle.

**Author Summary:** The centrosomes are responsible for organizing the mitotic spindle a microtubule-based structure that centers, then segregates, the chromosomes during cell division. When a cell divides it normally possesses two centrosomes, allowing it to build a bipolar spindle and accurately segregate the chromosomes to two daughter cells. Appropriate control of centrosome number is therefore crucial to maintaining genome stability. Centrosome number is largely controlled by their regulated duplication. In particular, the protein Plk4, which is essential for duplication, must be strictly limited as an overabundance leads to excess centrosome duplication. We have identified protein phosphatase 1 as a critical regulator of the *C. elegans* Plk4 homolog (known as ZYG-1). When protein phosphatase 1 is down-regulated, ZYG-1 levels increase leading to centrosome amplification. Thus our work identifies a novel mechanism that limits centrosome duplication.

## Introduction

In mitotic cells the centrosome serves as the primary microtubule-organizing center and consists of two centrioles surrounded by a proteinaceous pericentriolar matrix (PCM). During mitosis the centrosomes organize the poles of the spindle, therefore maintaining appropriate centrosome numbers promotes spindle bipolarity and faithful chromosome segregation. Regulated centrosome duplication is the primary mechanism by which centrosome number is controlled, and involves building a new daughter centriole adjacent to each pre-existing mother centriole. Two features of centriole duplication maintain appropriate centrosome numbers: first, centriole duplication is limited to occurring only once per cell cycle. Second, only a single daughter centriole is assembled in association with each pre-existing mother centriole.

A conserved set of five centriole duplications factors, SPD-2/CEP192, ZYG-1/Plk4, SAS-6, SAS-5/STIL/Ana2, are required for daughter centriole assembly and their individual loss results in centriole duplication failure (reviewed in [1]). Conversely, individual over expression of a subset of these duplication factors, Plk4, SAS-6 and STIL/SAS-5, leads to centriole over-duplication (the production of more than a single daughter) leading to a condition known as centriole amplification (the accumulation of an excess number of centrioles) [2–6]. Interestingly, the three factors whose overexpression leads to centriole amplification have been identified as key players in the initial steps of centriole duplication. In human cells Plk4 phosphorylates STIL to trigger centriolar recruitment of SAS-6, which initiates formation of the cartwheel, the central scaffolding structure of the new centriole [7–12]. Similarly in *C. elegans* the Plk4 homolog ZYG-1 recruits a complex of SAS-5 and SAS-6 through direct physical association with SAS-6 to initiate centriole duplication [13,14].

Because Plk4 overexpression causes the formation of extra daughter centrioles, Plk4 protein levels must be tightly regulated *in vivo.* One mechanism that regulates Plk4 levels is its SCF-mediated degradation promoted by autophosphorylation, whereby the active kinase induces its own destruction [15–17]. Degradation of Plk4 homologs in *Drosophila,* (plk4/Sak) and *C. elegans* (ZYG-1) are similarly regulated by their SCF-mediated targeting for degradation [18,19]. In addition, recent studies have shed light on temporal and spatial regulation of Plk4 levels. Plk4 initially localizes in a broad ring around the mother centriole until, coincident with the initiation of duplication, it becomes restricted to a small focus marking the location of daughter centriole assembly [8,20,21]. Emerging evidence suggests that this transition, which seems to be a key step in ensuring only a single daughter centriole is assembled, relies on spatially regulated Plk4 degradation [8,22]. Because STIL can both activate Plk4 [10,22,23], and also protect it from degradation [22] it is proposed that centriolar recruitment of a small focus of STIL at the G1/S transition triggers broad Plk4 activation and degradation via autophosphorylation, while protecting a local focus. Thus STIL limits the centriolar distribution of Plk4, promoting the assembly of a single daughter centriole. These studies reveal the central role that regulated destruction of ZYG-1/Plk4 plays in controlling centriole number, and highlight the importance of better understanding how the stability of Plk4 is regulated.

Protein phosphatase 1 (PP1) is a major cellular phosphatase that plays well-characterized roles in diverse processes including glycogen metabolism, circadian rhythms and cell division. Humans possess a single PP1α gene, a single PP1β gene (also known as PP1δ) and a single PP1γ gene. The different isoforms are >85% identical and show largely overlapping roles, although some functional specialization has been identified [24]. *C. elegans* has four PP1 catalytic subunits: a single broadly-expressed PP1β homolog, GSP-1, and three PP1α homologs, GSP-2, which is also widely expressed, and GSP-3 and GSP-4, whose expression is limited to the male germ line [25,26]. The core catalytic subunit of PP1 can directly bind a subset of substrates, but functional specificity is largely conferred by its interaction with regulatory proteins. Over 200 PP1 interactors exist, which regulate PP1 through modulating substrate specificity, enzyme localization or by inhibition or activation of phosphatase activity [27]. Two evolutionarily conserved PP1 regulators that play a role in cell division are inhibitor 2 (I-2) and SDS22. I-2 was originally identified as an inhibitor of PP1 and shows potent inhibitory activity *in vitro,* although interestingly the yeast homolog of I-2, GLC8, can also stimulate PP1 activity [28–30]. Down regulation of I-2 in *Drosophila* or human cells leads to chromosome mis-segregation, which is proposed to result from mis-regulation of Aurora B [31,32]. Similarly SDS22 antagonizes Aurora B autophosphorylation, downregulating Aurora B kinase activity [33]. Although it has been noted that PP1α regulates centrosome cohesion, [34,35], no role for PP1 in regulating centrosome duplication has previously been found.

Here we report a novel PP1-dependent pathway that plays a critical role in ensuring that each mother centriole produces one and only one daughter centriole during each cell cycle. We show that PP1 together with two of the most conserved PP1 regulators, I-2 and SDS-22 downregulates ZYG-1 protein abundance. Our data indicate that PP1 regulates ZYG-1 levels post-translationally and we demonstrate that loss of PP1-mediated regulation leads to ZYG-1 overexpression and centriole amplification through the production of multiple daughter centrioles.

## Results

### Loss of I-2^szy-2^, a conserved PP1 regulator, rescues the centriole assembly defect of zyg-1(it25) embryos

ZYG-1 is essential for centriole assembly: when hermaphrodites carrying the temperature sensitive *zyg-1(it25)* mutation are grown at the non-permissive temperature of 24°C centriole duplication fails, resulting in 100% embryonic lethality. To identify additional regulators of centriole assembly, we screened for suppressors of the *zyg-1(it25)* phenotype and isolated mutations in 20 *szy* (<u>s</u>uppressor of *<u>zy</u>g-1)* genes that rescue embryonic survival [36]. We mapped one of these mutations, *szy-2(bs4),* to the Y32H12A.4 locus on chromosome III, hereafter referred to as *szy-2.* The *szy-2(bs4)* mutation is a single base pair change (G to A) in a splice donor site, which is predicted to alter splicing of the *szy-2* transcript resulting in a frame-shift (Fig 1A). Analysis of the SZY-2 sequence shows that it is homologous to inhibitor-2 (I-2), a conserved regulator of protein phosphatase 1 [30]. Using antibodies raised against I-2^SZY-2^ we confirmed that the *szy-2(bs4)* mutation significantly reduces I-2 ^SZY-2^ protein levels, although residual I-2^SZY-2^ is still detected, indicating that at least some message is properly spliced in the mutant (Fig 1B). At 24°C the embryonic viability of *zyg-1(it25); szy-2(bs4)* double mutant embryos is 83%, significantly higher than *zyg-1(it25)* (0% viability). To determine if suppression of the embryonic lethal phenotype was associated with restoration of centriole duplication, we imaged *zyg-1(it25)* and *zyg-1(it25); szy-2(bs4)* embryos expressing GFP::tubulin and mCherry::histone (Fig 1C). Normally at fertilization the sperm delivers a single pair of centrioles to the acentrosomal egg, and during the first cell cycle the centrioles separate, duplicate and form the poles of the mitotic spindle (Fig 1C (top panel), Fig S1A and movie S1). Subsequent duplication events ensure the formation of bipolar spindles in later divisions. When the first round of centriole duplication fails, as in *zyg-1(it25)* mutants, the two sperm-derived centrioles still separate and organize the poles of the first mitotic spindle leading to a normal first division. However as each daughter cell inherits a single centriole, monopolar spindles assemble in the 2-cell embryo (Fig 1C (middle panel), Fig S1B and movie S2). As this phenotype is indicative of centriole duplication failure, we scored the presence of monopolar spindles in 2-cell stage embryos and found that >80% of centrioles duplicated during the first round of duplication in *zyg-1(it25);szy-2(bs4)* double mutant embryos (Figs 1C (bottom panel), 1D & movie S3). In contrast, this first duplication event always failed in *zyg-1(it25)* control embryos (Fig 1C (middle panel) & D). We confirmed that this effect is specific by using RNAi to deplete SZY-2 in *zyg-1(it25)* worms; RNAi of *szy-2* but not the non-essential gene *smd-1* (control RNAi), restored centriole duplication to the *zyg-1(it25)* mutant (Fig 1D). Moreover the deletion allele *szy-2(tm3972),* in which the C-terminal 190 residues are removed, was also able to suppress the *zyg-1(it25)* phenotype (Fig 1D). Together these data demonstrate that reducing I-2^SZY-2^ function suppresses *zyg-1(it25)* and suggest that modulating PP1 activity may impact centriole duplication.

**Figure 1.**
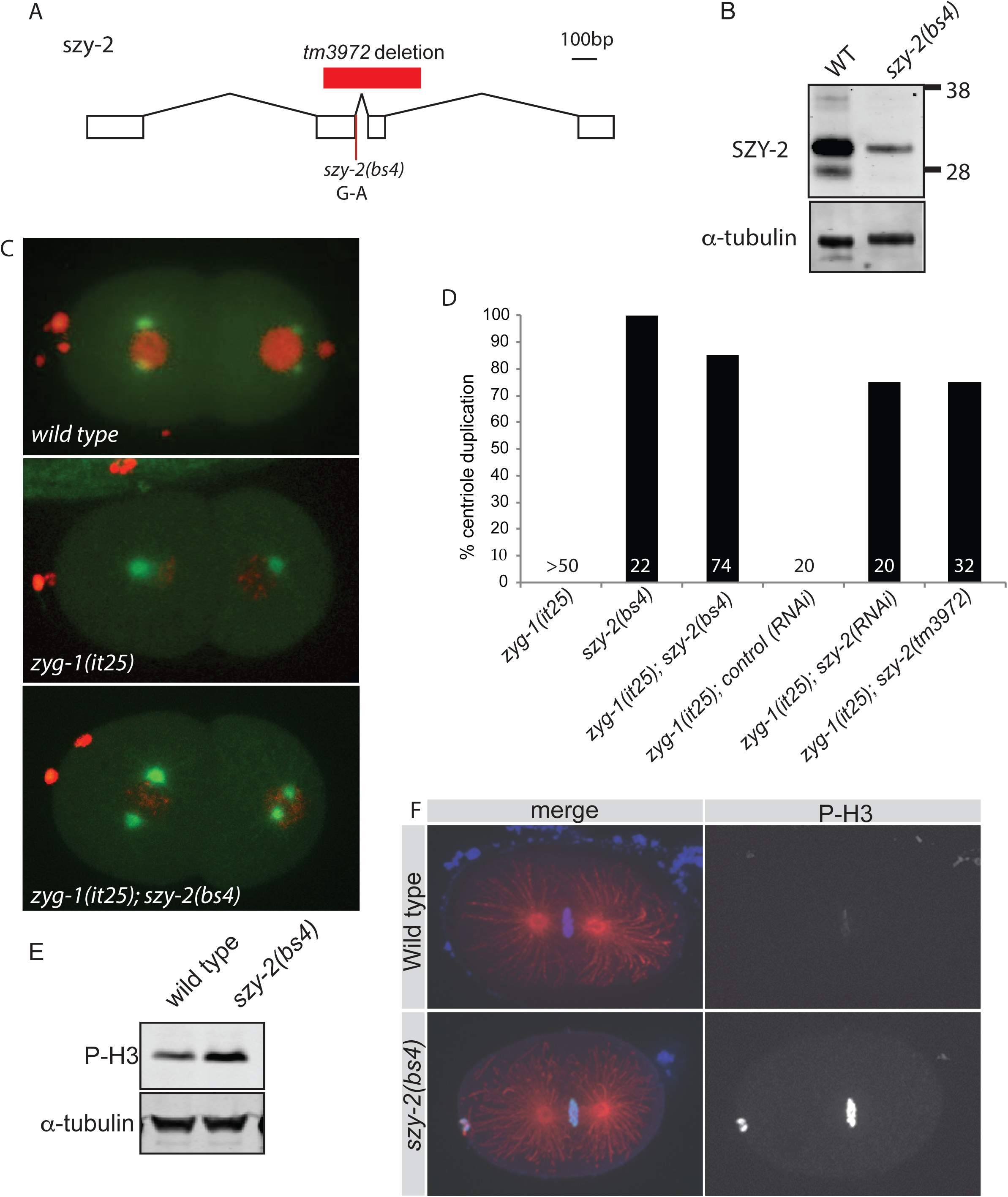
Loss-of-function mutations in the PP1 regulator I-2^SZY-2^ rescues the *zyg-1(it25)* phenotype. A) Scale diagram of the structure of the *szy-2* gene, indicating the location of the *tm3972* deletion and the *szy-2(bs4)* splice site mutation. B) Western blot of embryo extracts from wild-type and *szy-2(bs4)* mutants showed a reduced level of the SZY-2 protein. C) Stills from movies of embryos expressing mCherry::H2B and GFP::tubulin grown at 24°C. Top, wild type; Middle, *zyg-1(it25)* mutant at the 2-cell stage following centriole duplication failure; bottom, *zyg-1(it25); szy-2(bs4)* double mutants at 2 cell-stage showing a rescue of the centriole duplication. D) Quantification of centriole duplication failure at 24°C in *zyg-1(it25)* mutants and in *zyg-1(it25)* mutants in which *szy-2* activity has been down-regulated by mutation or RNAi. Number of centriole duplication events analyzed is indicated. E) Western blot of embryo extracts, showing levels of phospho-histone H3 in wild type and *szy-2(bs4)* embryos. F) Phospho-histone H3 staining of wild-type and *szy-2(bs4)* embryos at first metaphase. Left, merge: DNA blue, Microtubules red, Phospho-histone H3 green. Right, phospho-histone H3.

### Loss of szy-2 decreases PP1 activity

Because *szy-2* encodes the *C. elegans* homolog of I-2, this suggested that the *szy-2(bs4)* allele may alter PP1 activity, resulting in the observed suppression of centriole duplication failure. I-2 is conserved from yeast through humans and has been described as both an inhibitor and an activator of PP1 [28–30]. Analyses *in vitro* have shown that *C. elegans* I-2^SZY-2^ can inhibit rabbit PP1 activity, but the *in vivo* role of *szy-2,* in particular in relation to endogenous worm PP1 subunits, has not previously been investigated [30]. We sought to determine whether the important function of I-2^SZY-2^ with respect to *zyg-1(it25)* suppression was as an inhibitor or an activator of PP1. To determine whether PP1 activity is up-or down-regulated in the *szy-2(bs4)* mutants we wanted to monitor the phosphorylation levels of a known PP1 substrate. One such substrate is histone H3, which is phosphorylated in the early stages of mitosis; this modification is removed by PP1 [37,38]. To investigate whether I-2^SZY-2^ is an activator or inhibitor of PP1 we analyzed phospho-histone levels in *szy-2(bs4)* mutant embryos. We used a phospho-histone H3 antibody to detect phosphorylated histone H3 levels in embryos and found them to be greatly elevated in *szy-2(bs4)* when compared with the wild type (Fig 1E & F), a result reminiscent of PP1 depletion (Fig 3A; [25,39]). Phospho-histone levels were elevated in both mixed stage embryos (Fig 1E) as well as during the metaphase stage of mitosis (Fig. 1F). This result indicates that the *szy-2(bs4)* mutation reduces PP1 activity and suggests that I-2^SZY-2^ normally acts as an activator of PP1. Consistent with I-2^SZY-2^ being a positive regulator of PP1, *szy-2(bs4)* mutant embryos exhibit a chromosome mis-segregation defect [36] that is similar to that of embryos depleted of PP1 (movie S4). There is precedent for I-2 acting an activator of PP1: in yeast I-2^GLC8^ binds and inactivates PP1^GLC7^, but upon phosphorylation of I-2^GLC8^, PP1 becomes activated [29]. The phosphorylated residue is conserved in all I-2 homologs including *C. elegans,* although it is not clear whether I-2^SZY-2^ is similarly regulated in the worm [30].

### Loss of a second PP1 regulator, sds-22, rescues the centriole duplication failure of zyg-1(it25) embryos

Because we had identified a mutation in I-2^szy-2^ as a suppressor of *zyg-1(it25),* we speculated that loss of additional PP1 regulatory proteins may have a similar effect. To test this hypothesis we identified seven *C. elegans* proteins which show a high degree of conservation with human PP1 regulators. We then used RNAi to deplete each factor in *zyg-1(it25)* worms, and monitored embryonic viability (Fig S2). RNAi of only the *sds-22* gene significantly increased embryonic viability of the *zyg-1(it25)* embryos. SDS-22 is a leucine-rich repeat (LRR) domain containing protein that localizes PP1 during mitosis, regulating mitotic progression [33,40–42]. Although *sds-22(RNAi)* had only a modest effect on *zyg-1(it25)* embryonic viability, direct observation of centriole duplication in *zyg-1(it25); sds-22(RNAi)* embryos, revealed 70% of centriole duplication events occurred normally (Fig 2B). The disparity between the strength of suppression of embryonic lethality and the strength of suppression of centriole duplication failure is likely explained by a combination of two factors. First, we find that on average, 30% of the duplication events in *zyg-1(it25); sds-22(RNAi)* embryos fail; such a high degree of cell division failure during early development would result in embryonic lethality. Second, since *sds-22* is an essential gene, RNAi knockdown may cause embryonic lethality even though centriole duplication is normal. Since reducing SDS-22 function is able to suppress the *zyg-1(it25)* phenotype we hypothesized that one of the suppressors of *zyg-1(it25)* recovered in our genetic screen[36] may have a mutation in *sds-22.* We therefore sequenced the *sds-22* coding region in those *szy* mutants that mapped close to the *sds-22* genetic locus. The *szy-6(bs9)* strain contained a G-to-A missense mutation in the coding region of *sds-22* which results in a single amino-acid substitution (G224S) within the eighth LRR (Fig 2A). To confirm that *bs9* was an allele of *sds-22* we performed a complementation test with *sds-22(tm5187)* a deletion allele that exhibits a fully-penetrant larval-lethal phenotype (Figs 2A & C). While the *szy-6(bs9)* homozygotes exhibited minimal embryonic lethality, the trans-heterozygotes *(bs9/ tm5187)* exhibited 100 percent embryonic lethality confirming that *bs9* and *tm5187* are allelic (Fig 2C). We have therefore renamed *the szy-6(bs9)* allele *sds-22(bs9).* Although by itself, the *sds-22(bs9)* mutation does not appear to affect centriole duplication, *zyg-1(it25); sds-22(bs9)* double mutant embryos exhibit an approximately 50% success rate of centriole duplication, comparable to levels seen after SDS-22 RNAi depletion (Fig 2B). Together these data show that reducing the function of either of two PP1 regulatory proteins, I-2^SZY-2^ or SDS-22, is able to partially suppress the failure of centriole duplication caused by the *zyg-1(it25)* mutation.

**Figure 2.**
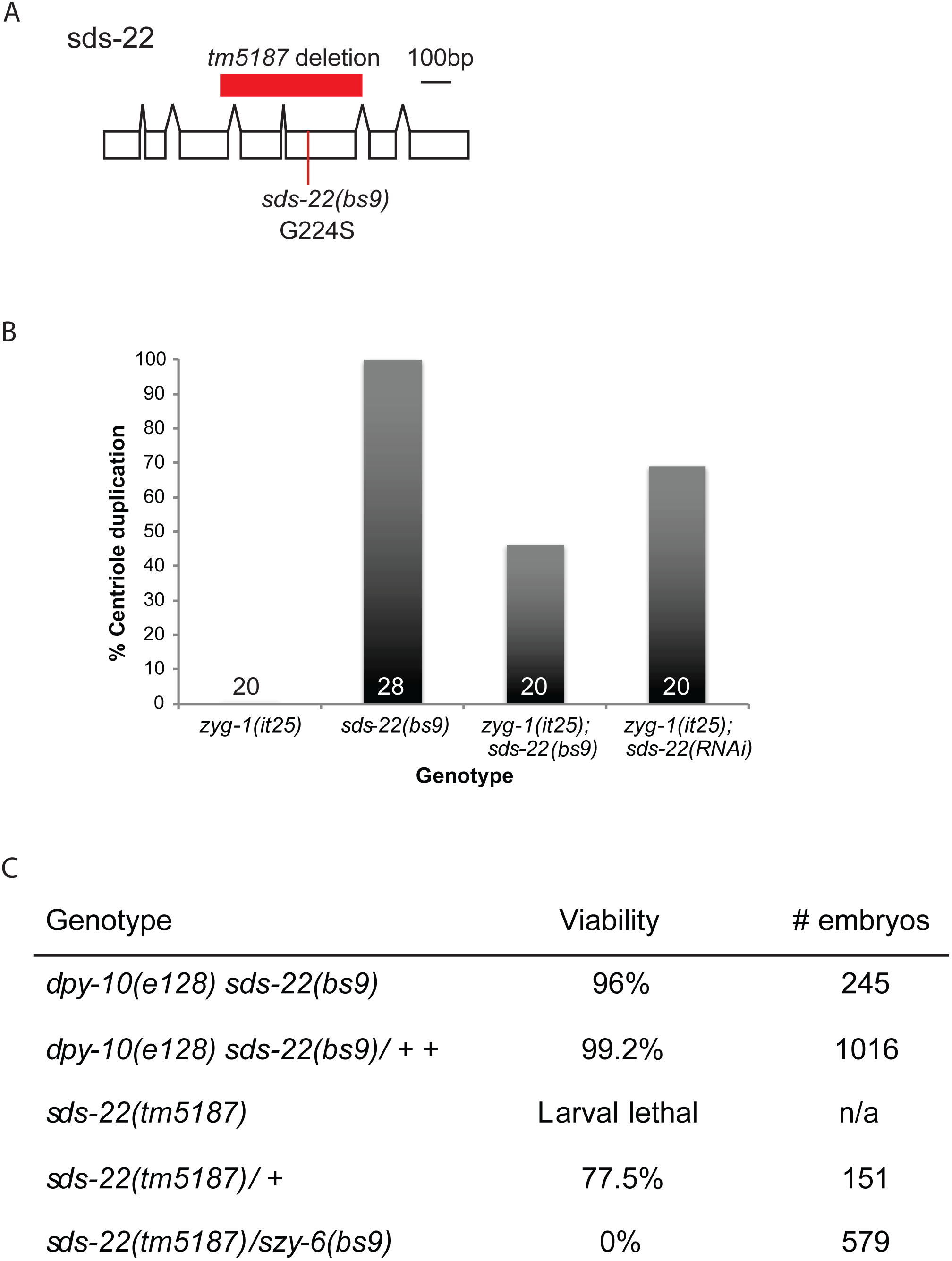
Reducing SDS-22 activity rescues the *zyg-1(it25)* phenotype. A) Scale diagram of the structure of the *sds-22* gene, indicating the location of the *tm5187* deletion and the *sds-22(bs9)* substitution. B) Quantification of centriole duplication when *zyg-1(it25)* is rescued by the *sds-22(bs9)* mutation or *sds-22(RNAi).* Number of centriole duplication events analyzed is indicated. C) Complementation test showing *bs9* and *tm5187* are allelic. Hermaphrodites of the indicated genotype were shifted to 25°C at the L4 stage and embryonic lethality was determined over the next 24 hours. The *szy-6(bs9)* allele was marked with the closely linked dpy-10(e128) mutation.

### Loss of PP1β^GSP-1^ activity rescues zyg-1(it25)

Our data suggest that I-2^SZY-2^ is an activator of PP1, and work in vertebrate cells has shown that the SDS-22 ortholog positively regulates PP1 activity [33]. Because the *szy-2(bs4)* and *sds-22(bs9)* mutations rescue *zyg-1(it25)* phenotypes it followed that directly lowering PP1 activity should also suppress the *zyg-1(it25)* phenotype. Embryos express two PP1 catalytic subunits encoded by the genes *gsp-1* and *gsp-2.* Similar to what is seen in the *szy-2(bs4)* mutant (Fig 1E & F), co-depletion of GSP-1 and GSP-2 by RNAi resulted in an increase in phospho-histone H3 staining of mitotic chromosomes (Fig 3A). We next tested whether individually reducing the activity of either PP1^GSP-1^ or PP1^gsp-2^, could suppress the centriole duplication failure seen in *zyg1(it25)* embryos. Although the *gsp-1(tm4378)* deletion allele is sterile, precluding its use in our analysis, we were able to specifically deplete PP1^GSP-1^ by RNAi (Fig 3B). When PP1^GSP-1^ was depleted in *zyg-1(it25)* embryos we saw a robust restoration of centriole duplication (90% of centrioles; Fig 3B, C, D & G). In contrast, the putative null allele *gsp-2(tm301)* [25] did not suppress the *zyg-1(it25)* phenotype and we never observed centriole duplication in the *zyg-1(it25); gsp-2(tm301)* double mutant embryos (Fig 3E, F & G). The *C. elegans* PP1α^GSP-2^ and PP1β^GSP-1^ isoforms show a high degree of identity and are both expressed in the early embryo [25]. Since depletion of only the PP1β^GSP-1^ isoform can suppress the *zyg-1(it25)* allele this suggests either 1) a divergence in function of the two enzymes, with only PP1β^GSP-1^ being involved in the regulation of centriole duplication, or 2) that, despite partial redundancy between the two enzymes, *zyg-1(it25)* mutants provides a sensitized background where loss of PP1β^GSP-1^ alone is sufficient to subvert normal controls. Indeed, two observations support the later possibility. First, we have only observed defects in chromosome segregation after co-depletion of both PP1β^GSP-1^ and PP1α^GSP-2^ suggestive of some functional redundancy (movie S4). Second, we show below that co-depletion of PP1β^GSP-1^ and PP1α^GSP-2^ disrupts the normal pattern of centriole duplication whereas single depletions of either phosphatase have no effect (Fig 5C, D, E & movie S5).

**Figure 3.**
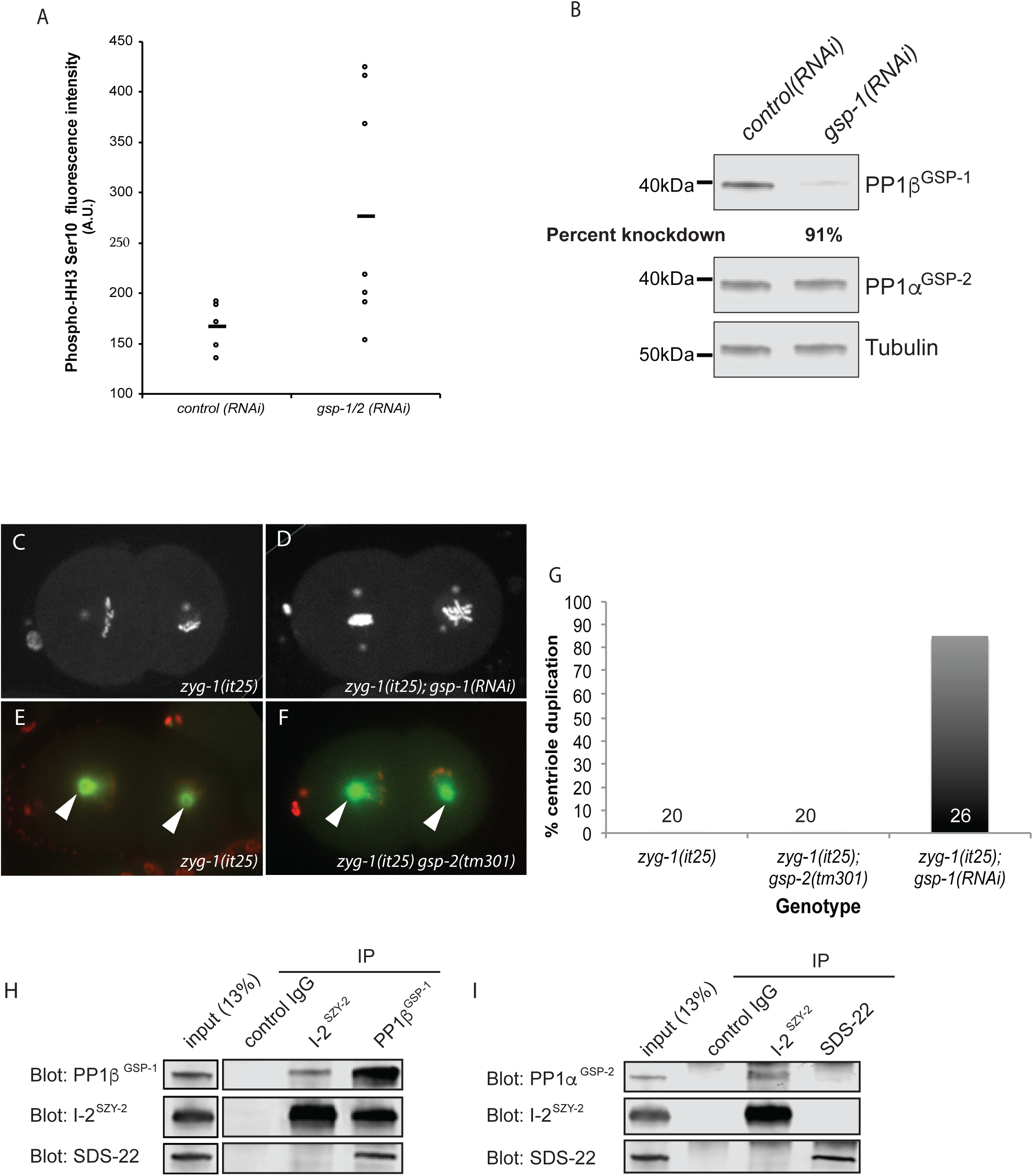
PP1 functions with I-2^SZY-2^ and SDS-22 to regulate centriole duplication. A) Comparison of phospho-HH3 levels in control and PP1β^GSP-1^ & PP1α^GSP-2^ co-depleted embryos as determined by quantitative immunofluorescence microscopy. The mean is indicated by the bar. B) Western blot demonstrating specificity of *gsp-1(RNAi).* C&D) Stills from time-lapse recordings of embryos of the indicated genotypes, expressing GFP::histone and GFP::SPD-2 grown at the normally restrictive temperature of 24°C. E&F) Stills from time-lapse recordings of embryos of the indicated genotypes, expressing mCherry::histone and GFP::tubulin grown at the normally restrictive temperature of 24°C; centrosomes are indicated by arrow heads. G) Quantitation of centriole duplication in indicated strains. Number of centriole duplication events scored is indicated. H) Western blots of immunoprecipitated material from whole worm extracts using control IgG, I-2^SZY-2^, or PP1β^GSP-1^ antibodies, and probed with I-2^S7Y-2^, PP1β^GSP-1^, or SDS-22 antibodies as indicated. I) Western blots of immunoprecipitated material from whole worm extracts using control IgG, I-2^S7Y-2^, or SDS-22 antibodies, and probed with I-2^S7Y-2^ or PP1α^GSP-2^, or SDS-22 antibodies.

Our data indicate that individually reducing the function of either I-2^SZY-2^, SDS-22 or PP1β^GSP-1^ can partially suppress the *zyg-1(it25)* centrosome duplication failure. Using GFP fusions we were, however, unable to detect centrosome localization of any of the proteins (Fig S3). Nevertheless we reasoned that all three proteins work in a common process and sought to determine whether they physically interact. When we immunoprecipitated I-2^SZY-2^ from *C. elegans* embryonic extracts, we found by immunoblotting that we could detect co-precipitated PP1β^GSP-1^ and reciprocally when we immunoprecipitated PP1β ^GSP-1^ we were able to pull down I-2 ^SZY-2^ (Fig 3H), confirming that *C. elegans* I-SZY-2 GSP-1 2 is a PP1 binding protein *in vivo.* Similarly, antibodies against PP1p ' co-precipitated SDS-22 (Fig 3H). However, we were unable to detect any interaction between I-2^SZY-2^ and SDS-22, suggesting that a complex simultaneously containing all three proteins does not form. Additional IP-western experiments demonstrated that I-2^SZY-2^ also interacts with PP1α^GSP-2^, but did not reveal an interaction between SDS-22 and PP1α^GSP-2^ (Fig 3I). To confirm and extend these results we also individually SZY-2 GSP-1 immunoprecipitated I-2^SZY-2^, SDS-22, and PP1β^GSP-1^ and analyzed the immunoprecipitated material by mass spectrometry (Fig S4). Using this approach we found that I-2^SZY-2^ and SDS-22 could independently interact with both PP1β^GSP-1^ and PP1α^GSP-2^ but not with each other (Fig S4). We conclude that components of the PP1-dependent regulatory pathway physically interact but that I-2^SZY-2^ and SDS-22 do not appear to reside in the same complex.

Finally, we have found that although both I-2^SZY-2^ and SDS-22 interact with PP1 and positively regulate PP1's function in controlling centriole duplication, the two regulators do not always function together to control PP1 activity. Specifically we have found that while loss of I-2^SZY-2^ activity results in an increase in the level of phospho-histone H3 in mitotic chromosomes (Fig 1F), no such increase is observed upon loss of SDS-22 activity (Fig S5B). Thus I-2^SZY-2^ appears to function independently of SDS-22 to mediate the role of PP1 in regulating histone H3 phosphorylation. Curiously, loss of SDS-22 resulted in an elevated phospho-histone H3 signal as detected by immunoblotting (Fig S5A). Because SDS-22 does not control the chromatin content of phospho-histone H3, we speculate that loss of SDS-22 leads to a mitotic delay; this would explain the elevated signal of phospho-histone H3 in mixed-stage worm extracts.

### Aurora B, Aurora A and Plk1 are not required for szy-2(bs4) suppression of zyg-1(it25)

Given that PP1 is a mitotic phosphatase that antagonizes many known mitotic kinases we wanted to investigate whether these relationships contribute to the regulation of centriole duplication. In *Drosophila* and human cells, I-2 is implicated in regulation of Aurora B and its depletion results in chromosome mis-segregation [31,32]. Similarly in humans, SDS22 antagonizes Aurora B autophosphorylation, downregulating Aurora B kinase activity [33]. PP1, I-2 and SDS22 therefore seem to cooperatively regulate chromosome segregation by modulating Aurora B activity. In *C. elegans* PP1 also regulates Aurora B, contributing to its correct localization during meiosis and to chromosome segregation in mitosis [39,43]. Although Aurora B has not been implicated in centriole duplication we wanted to determine whether *szy-2(bs4)* suppresses centriole duplication failure through upregulation of Aurora B activity. We therefore RNAi-depleted Aurora B^AIR-2^ in *szy-2(bs4); zyg-1(it25)* worms and monitored centriole duplication (Fig S6A, B & C). Although we observed failures in chromosome segregation and cytokinesis consistent with successful Aurora B depletion (Fig S6C;[44]), centriole duplication was unperturbed (Fig S6A), indicating that *szy-2(bs4)* does not regulate centriole duplication by modulating Aurora B activity.

Since PP1 and I-2 are associated with regulation of Aurora A activity [45] and Aurora A plays a conserved role in centrosome separation and maturation [46,47] we tested whether Aurora a^air-1^ activity is required for suppression of *zyg-1 (it25)* by *szy-2(bs4).* After exposing double mutants to *air-1(RNAi)* we observed reduced SPD-2 localization and incomplete centrosome separation, consistent with loss of Aurora A activity (Fig S7H), however we did not see a perturbation of centriole duplication (Fig S6A, Fa & Fb), suggesting that Aurora A is not required for suppression of *zyg-1(it25)* by *szy-2(bs4).* In vertebrate cells PP1 also has a recognized role in antagonizing Plk1 [48], a kinase with an established role in centriole duplication [49]. However, depletion of *plk-1* in *zyg-1(it25), szy-2(bs4)* embryos did not affect centriole duplication (Fig S6A) even though PLK-1 activity was clearly compromised (Fig S6D). In summary our data suggest that reducing PP1 activity does not restore centriole duplication in the *zyg-1(it25)* mutant by relieving antagonism, and thus increasing the relative activity, of Aurora B, Plk1 or Aurora A, which are known PP1 antagonists.

### I-2^szy-2^ controls SPD-2 levels at the centrosome

During the course of our analyses we noted that the level of the coiled-coil protein SPD-2 was elevated at the centrosome in *szy-2(bs4)* mutant embryos. SPD-2 is a component of both the PCM and centrioles and is required for centriole duplication and PCM assembly [50,51]. We quantified the effect on SPD-2 levels in *szy-2(bs4)* embryos expressing SPD-2::GFP and found a 1.5-fold increase in centrosomal SPD-2 levels throughout the cell cycle (Fig S7A). When we compared levels of endogenous SPD-2 at the centrosome, we found a similar increase in centrosome-associated SPD-2 in *szy-2(bs4)* embryos (Fig S7B-D). Since SPD-2 plays a positive role in centriole duplication, we reasoned that its increased levels at the centrosome in the *szy-2(bs4)* mutant might be responsible for PP1-mediated suppression. To test this possibility, we utilized a codon-optimized *spd-2::gfp* transgene [52] to overexpress SPD-2 protein in the *zyg-1(it25)* mutant. Despite a large increase in centrosome-localized SPD-2 (Fig S7E & F) we did not see any suppression of *zyg-1(it25)* embryonic lethality (Fig S7G), indicating that centriole duplication failure persisted. Thus by itself, a general elevation of the level of SPD-2 at the centrosome is not sufficient for suppression of *zyg-1(it25).* Conversely, we also find that the elevated level of centrosome-associated SPD-2 is not required for *szy-2(bs4)-mediated* suppression: Aurora A^AIR-1^ is required for SPD-2 recruitment to the centrosome [50], and depletion of aurora A^AIR-1^ in the *zyg-1(it25); szy-2(bs4)* double mutant drastically reduced the amount of SPD-2 at the centrosome, (Fig S7H) nevertheless, centriole duplication was not perturbed (Fig S6A, E & F). Thus, I-2^szy-2^ regulates both centriole duplication and PCM assembly, but the elevated SPD-2 levels at the centrosome observed in *szy-2(bs4)* mutants do not constitute the primary mechanism by which *szy-2(bs4)* suppresses the centriole duplication defect of *zyg-1(it25)* mutants.

### Reducing PP1 activity elevates ZYG-1 protein levels

Previous work has shown that the failure of centriole duplication in *zyg-1(it25)* mutants can be rescued by increasing centriole-associated levels of the mutant ZYG-1 protein [19,53]. We therefore sought to determine if decreasing PP1 activity affects ZYG-1 protein levels or localization. To determine whether the *szy-2(bs4)* mutation affects ZYG-1 abundance, we measured total ZYG-1 levels in early mixed-stage embryos using quantitative immunoblotting. Strikingly, in comparison to the wild type, *szy-2(bs4)* embryos consistently possessed approximately four-fold more ZYG-1 at the restrictive temperature of 25°C (Fig 4A & B). We then used quantitative immunofluorescence to see if the elevated level of total ZYG-1 protein resulted in increased levels of ZYG-1 at centrioles. As expected, the overall increase in ZYG-1 levels in embryos was associated with elevated ZYG-1 at the centrioles (Fig 4C, D & E). This increase however was cell cycle stage specific with increases observed in prophase and metaphase but not anaphase, of the first cell cycle (Fig 4C). We further sought to determine whether decreasing SDS-22 or PP1 similarly increased ZYG-1 levels. Quantification of centrosome-associated ZYG-1 in prophase revealed elevated levels of ZYG-1 in the *sds-22(bs9/tm5187)* trans-heterozygous mutant but a minimal increase after RNAi-depletion of PP1β^GSP-1^ (Fig 4D & E). Similarly depletion of PP1α^GSP-2^ had little effect on ZYG-1 levels at centrioles (Fig 4D & E). Since we only observed chromosome segregation defects when we depleted both PP1β^GSP-1^ and PP1α^GSP-2^ (movie S4), we wondered whether they also acted redundantly to regulate ZYG-1. We therefore co-depleted PP1β^GSP-1^ and PP1α^GSP-2^ and found an increase in centrosome-associated ZYG-1, consistent with the existence of redundancy between the two PP1 isoforms (Fig 4D & E). Overall our results suggest that reducing PP1 activity leads to an elevation of both total ZYG-1 levels, and of centriole-associated ZYG-1, providing a likely mechanism for the suppression of the *zyg-1(it25)* centriole duplication defect. Notably, this effect, of reduced PP1 activity leading to increased ZYG-1 levels, seems to be specific as we did not see a similar increase in SAS-6 levels in *szy-2(bs4)* embryos (Fig S8A, B & C). Finally, because ZYG-1 has also been implicated in positively regulating centrosome size [53], elevated ZYG-1 levels may also account for the observed increase in centrosomal SPD-2.

**Figure 4.**
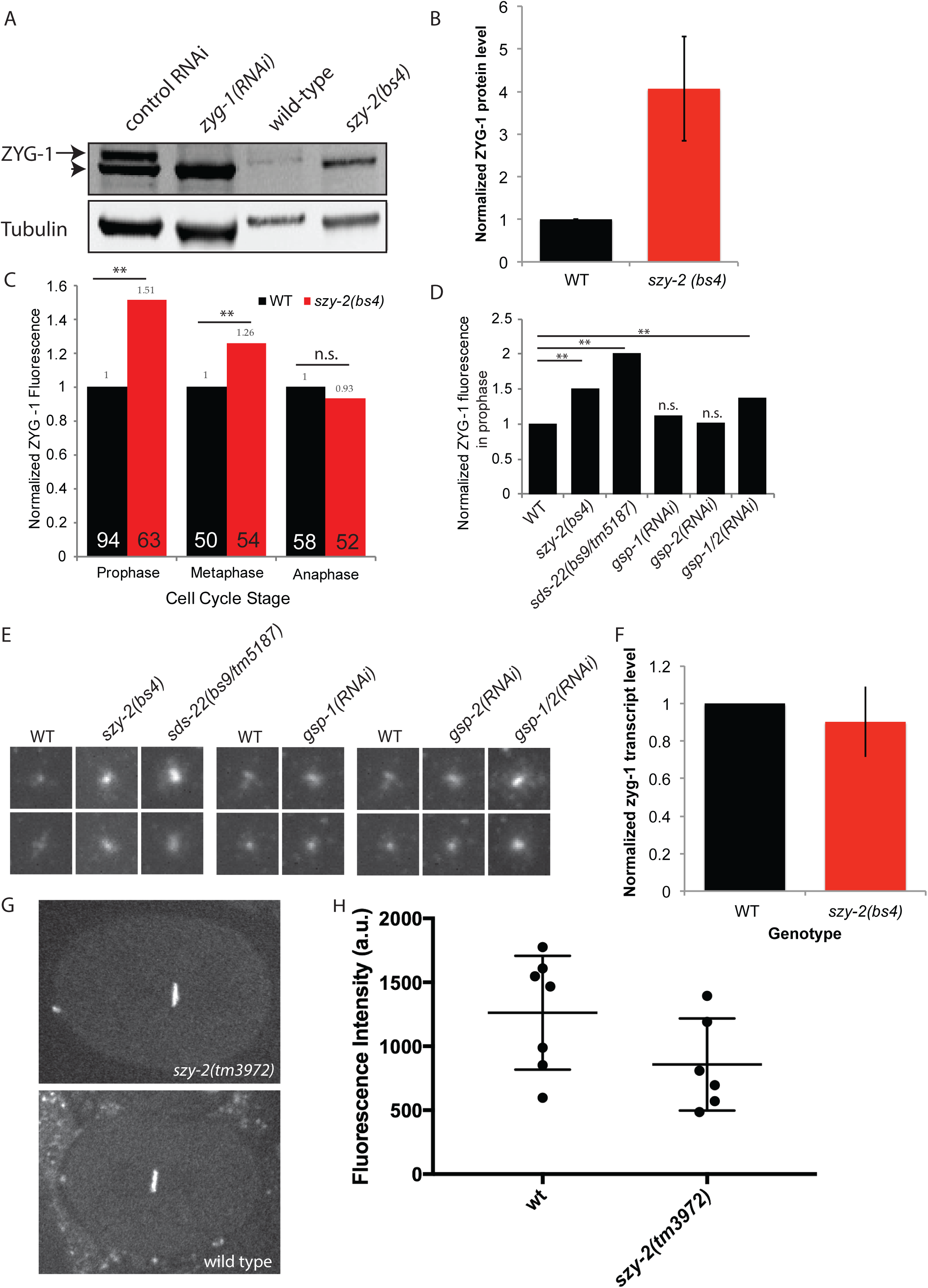
ZYG-1 protein levels are elevated in the *szy-2(bs4)* mutant. A) Western blot of embryo extracts from wild type and *szy-2(bs4)* embryos probed with a ZYG-1 antibody. The ZYG-1 band (arrow) is identified by its absence in *zyg-1(RNAi)* extracts. Non-specific proteins (arrowhead) are also bound by the ZYG-1 antibody and probably represent contaminants from the *E. coli* strain used for RNAi feeding experiments. B) Normalized ZYG-1 protein levels from two independent experiments. Error bars indicate standard deviation. C) Relative levels of centriole-localized ZYG-1 in wild-type and *szy-2(bs4)* one-cell embryos made by quantitative immunostaining. Levels at each stage are normalized to the wild-type control. Number of centrosomes analyzed indicated inside each bar. **p<0.01; n.s. no significant difference (Student's t-test). D) Centriole-localized ZYG-1 protein levels for embryos of indicated genotypes at first mitotic prophase. Levels normalized to the wild-type control. Number of centrosomes analyzed indicated inside each bar. **p<0.01; n.s. no significant difference from WT (Student's t-test). E) Representative images of centrosomal ZYG-1 levels in prophase. Centrosomes from 1-cell stage embryos of the indicated genotypes are grouped with a wild type sample from the same experiment. F) Quantification of *zyg-1* transcript levels in embryos by qRT-PCR. Shown is the average of two independent experiments. Error bars indicate standard deviation. G) Representative images of wild-type and *szy-2(tm3972)* embryos expressing GFP::histone driven by the *zyg-1* promoter and 3'-UTR. H) Quantitation of GFP intensities with mean and standard deviations indicated. No difference between stains was found (Student's t-test p>0.1).

### PP1 regulates ZYG-1 levels post-translationally

We next sought to determine how ZYG-1 levels are regulated by PP1. First, we measured *zyg-1* transcript levels by quantitative RT-PCR and found that wild-type and *szy-2(bs4)* embryos possessed similar levels of *zyg-1* mRNA (Fig 4F). The observed increase in ZYG-1 protein levels was not, therefore, due to an effect on transcription or mRNA stability. To test whether decreasing PP1 activity resulted in an effect on ZYG-1 translation efficiency, we crossed the *szy-2(tm3972)* allele into a strain carrying a GFP::histone reporter expressed under the control of the *zyg-1* regulatory sequences (promoter and UTRs) [54]. Since this reporter contains *zyg-1* regulatory sequences, but lacks the *zyg-1* coding sequence, it allows us to determine whether elevated ZYG-1 protein levels stem from alterations at the level of gene expression (transcription/translation) or from post-translational controls. Quantitative fluorescence intensity measurements of GFP::histone revealed that the *szy-2(tm3972)* mutation did not increase expression of the reporter relative to that observed in the wildtype strain (Fig 4G & H; Student's t-test p>0.1), indicating that reduced PP1 activity does not increase *zyg-1* translation via the 3‘-UTR. In order to determine whether PP1 might directly regulate ZYG-1 levels we tested for an interaction between the two proteins. Immunoprecipitation of I-2^szy-2^, SDS-22, and PP1β^GSP-1^ from worm extracts, followed by western blotting or mass spectrometry failed to detect an interaction between any of the three proteins and ZYG-1. Because ZYG-1 is a low abundance protein [14] and interactions between PP1 and its substrates may be transient we cannot however rule out a direct interaction between ZYG-1 and PP1. Cumulatively, therefore, our data point to PP1 regulating ZYG-1 protein levels either directly or indirectly via a post-translational mechanism.

### Reduction of PP1 activity causes centriole amplification

Elevated levels of the human and *Drosophila* homologs of ZYG-1 are associated with centriole amplification [2,5,6]. Although centriole amplification has not previously been observed in *C. elegans* embryos, given the increases in ZYG-1 levels we observed in the *szy-2(bs4)* and *sds-22(bs9/tm5187)* mutants we were intrigued whether this would be sufficient to subvert the normal regulation of centriole duplication, resulting in supernumerary centrosomes. We therefore performed time-lapse confocal microscopy of embryos expressing GFP::SPD-2 and mCherry::histone. Initial inspection of *szy-2* and *sds-22* mutants did not reveal any obvious defects during the first two cell cycles; centriole duplication proceeded normally and bipolar spindles assembled in all cells. Unexpectedly however, in embryos strongly impaired for either I-2^SZY-2^ or SDS-22 function, extra centrosomes were frequently observed at the four-cell stage (Fig 5A, B & movie S6). Closer inspection of the movies indicated that the extra centrosomes arose approximately synchronously from the spindle poles as the PCM dispersed near the completion of the second mitotic divisions (movies S6 & S7). In a single case (out of 79 analyzed) we were not able to trace the origin of one of these centrosomes to a spindle pole. Thus, it is possible that very infrequently in this mutant a centriole arises spontaneously in the cytoplasm via a *de novo* pathway. The production of extra centrosomes was most prevalent in *sds-22(bs9/tm5187)* trans-heterozygotes where nearly 40% of the spindle poles gave rise to more than two centrosomes (Fig 5B). Although less frequent, an identical defect was observed in *szy-2(tm3972)* embryos where supernumerary centrosomes arose with the same timing and spatial pattern as those observed in the *sds-22* mutant (Fig 5B). These results suggest that extra centrioles form during the second round of centriole duplication (which occurs in the 2-cell embryo) such that the extra centrioles only become apparent by confocal microscopy when mother and daughter centrioles separate as cells enter the third cell cycle (Fig S1C).

**Figure 5.**
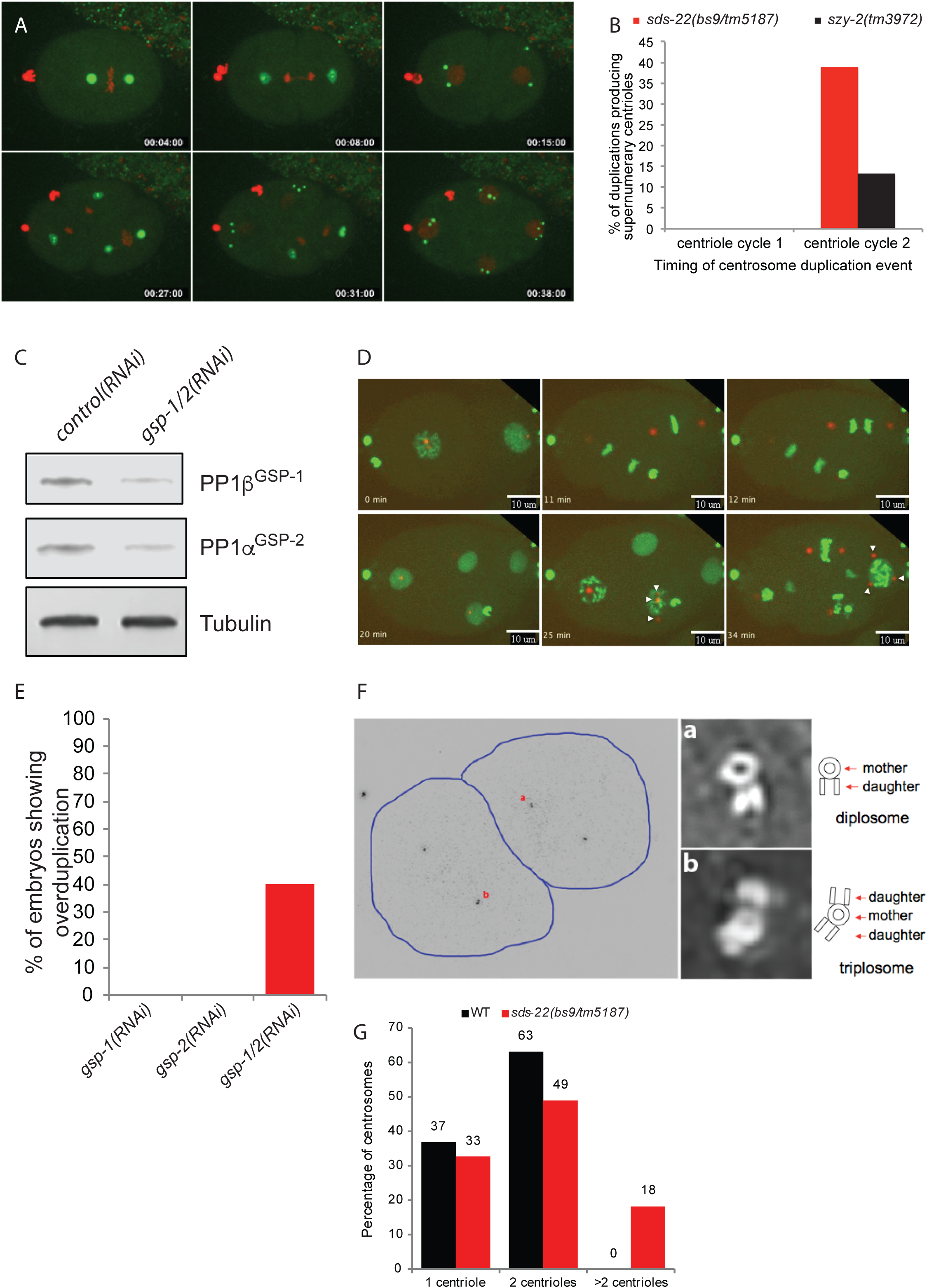
Decreasing PP1 activity leads to centriole overduplication. A) Frames from a time-lapse recording of an *sds-22(bs9/tm5187)* trans-heterozygous embryo expressing GFP::SPD-2 and mCherry::histone. Supernumerary centrosomes appear at the 4-cell stage. B) Quantification of centrosome over-duplication in *sds-22(bs9/tm5187)* and *szy-2(3972)* embryos during the first and second rounds of centriole duplication. For *sds-22(bs9/tm5187),* n=31/77 (first round/second round) and for *szy-2(3972),* n= 42/68 (first round/second round) C) Immunoblot showing extent of knockdown of PP1β^GSP-1^ and PP1α^GSP-2^ in *gsp-1; gsp-2* double RNAi embryos. D) Selected frames from a time-lapse recording of a *gsp-1(RNAi); gsp-2(RNAi)* embryo expressing GFP::histone and mCherry::SPD-2. Arrowheads indicate extra centrosomes. E) Quantitation of the frequency of supernumerary centrioles in embryos depleted of the indicated PP1 genes. F) Structured illumination microscopy (SIM) of a late 2-cell stage *sds-22(bs9/tm5187)* trans-heterozygous embryo reveals the presence of excess centrioles. The low magnification image on the left shows the positions of the centrioles stained for the centriole marker SAS-4. Images on the right are enlargements of the indicated centrosomes. G) Quantification of SIM data to indicate the number of centrioles observed per centrosome comparing wild type and *sds-22(bs9/tm5187)* transheterozygous mutants.

We next followed the fate of 79 centrosomes in *sds-22(bs9/tm5187)* trans-heterozygotes and found that all eventually accumulated PCM and participated in spindle assembly, often giving rise to multipolar spindles (movie S7). A minority (4/79 centrosomes), however, exhibited a one-cell-cycle delay before accumulating normal levels of PCM and participating in spindle assembly (Fig S9 and movie S7). This is somewhat reminiscent of the reversible "inactivation" of centrioles observed in *Drosophila* embryos following over-expression of Plk-4 [55]. The production of excess centrosomes continued in the ensuing cell cycles. We quantified the rate of over-duplication following the third centriole cycle of *sds-22(bs9/tm5187)* embryos and found a similar frequency of over-duplication (35%, n= 96 centrosomes). Thus our results suggest that over-duplication begins during the second centriole duplication event and continues through embryonic development.

To confirm that the excess centrosomes observed in *szy-2* and *sds-22* mutant embryos arose due to a reduction of PP1 activity, we also followed centrosomes during the first several cell cycles of embryos depleted for one or both PP1 catalytic subunits. Surprisingly, neither depletion of PP1β^GSP-1^ (n=10 embryos) nor of PP1α^GSP-2^ (n=11 embryos) resulted in the appearance of extra centrosomes during the first three cell cycles (Fig 5E). We therefore co-depleted both catalytic subunits (Fig. 5C) and found that excess centrosomes appeared at the four-cell stage in 4/10 embryos (Fig 5D, E & movie S5). Further, these extra centrosomes could function as microtubule-organizing centers and directed the formation of tripolar spindles (movie S8). Interestingly, we also observed occasional anaphase chromatin bridges in these embryos (movie S4), whereas in singly depleted embryos chromosomes always segregated normally. We conclude that PP1β^GSP-1^ and PP1α^GSP-2^ exhibit some level of functional redundancy in their roles in chromosome segregation and centriole duplication.

To further investigate the origin of the excess centrosomes we analyzed *sds-22(bs9/tm5187)* embryos by structured illumination microscopy (SIM), which has proven an effective means to observe centriole arrangement and number within the centrosome [56,57]. SIM imaging of SAS-4-stained embryos allowed us to resolve the basic structure of worm centrioles and we could clearly detect the normal arrangement of mother and daughter centrioles (Fig 5Fa). Strikingly, in *sds-22(bs9/tm5187)* embryos we were also able to find mother centrioles bearing more than one daughter (Fig 5Fb). We first detected extra daughter centrioles at the late 2-cell stage, consistent with the appearance of excess centrosomes at the four-cell stage. Furthermore, the appearance of extra daughter centrioles in association with a single mother is reminiscent of what is seen after Plk4 overexpression [2] and indicates that the excess centrosomes we observe in the 4-cell embryo originate from the formation of extra daughters during centriole duplication. Quantification of our SIM data reveals that while wildtype embryos never exceed two centrioles per centrosome, 18% of *sds-22(bs9/tm5187)* centrosomes contain more than 2 centrioles (Fig 5G). We did not observe any other unusual centriole configurations such as mother-daughter-granddaughter arrangements indicative of reduplication of centrioles during a single cell cycle. In summary, our data indicate that loss of PP1 activity results in overexpression of ZYG-1 and consequently centriole amplification. Amplification appears to be largely, if not entirely, driven by the production of multiple daughter centrioles. However, we cannot rule out that centriole reduplication and *de novo* formation also contribute to the excess centrioles observed after PP1 inhibition. In conclusion, we have described a new PP1-dependent mechanism that limits centriole duplication so that each mother produces one and only one daughter per cell cycle.

## Discussion

We have shown that PP1 activity is an important regulator of ZYG-1 levels in the *C. elegans* embryo and therefore that it is a critical regulator of centriole duplication. Although PP1α has previously been implicated in the regulation of centrosome separation at the beginning of mitosis [34], this is the first indication that PP1 regulates centriole duplication. Our data indicate that PP1 inhibits centriole duplication by restraining the accumulation of ZYG-1 in the embryo either directly or indirectly (Fig 6). Previous work has shown that Plk4 levels can be regulated by SCF-mediated degradation promoted by autophosphorylation [15–17]. Similarly in *C. elegans* the SCF complex is involved in regulating ZYG-1 levels and depletion of SCF components leads to an increase in centrosome-associated ZYG-1 [19]. Our finding that down-regulation of PP1 activity leads to centrosome amplification implicates PP1 as an additional regulator of ZYG-1 levels.

**Figure 6.**
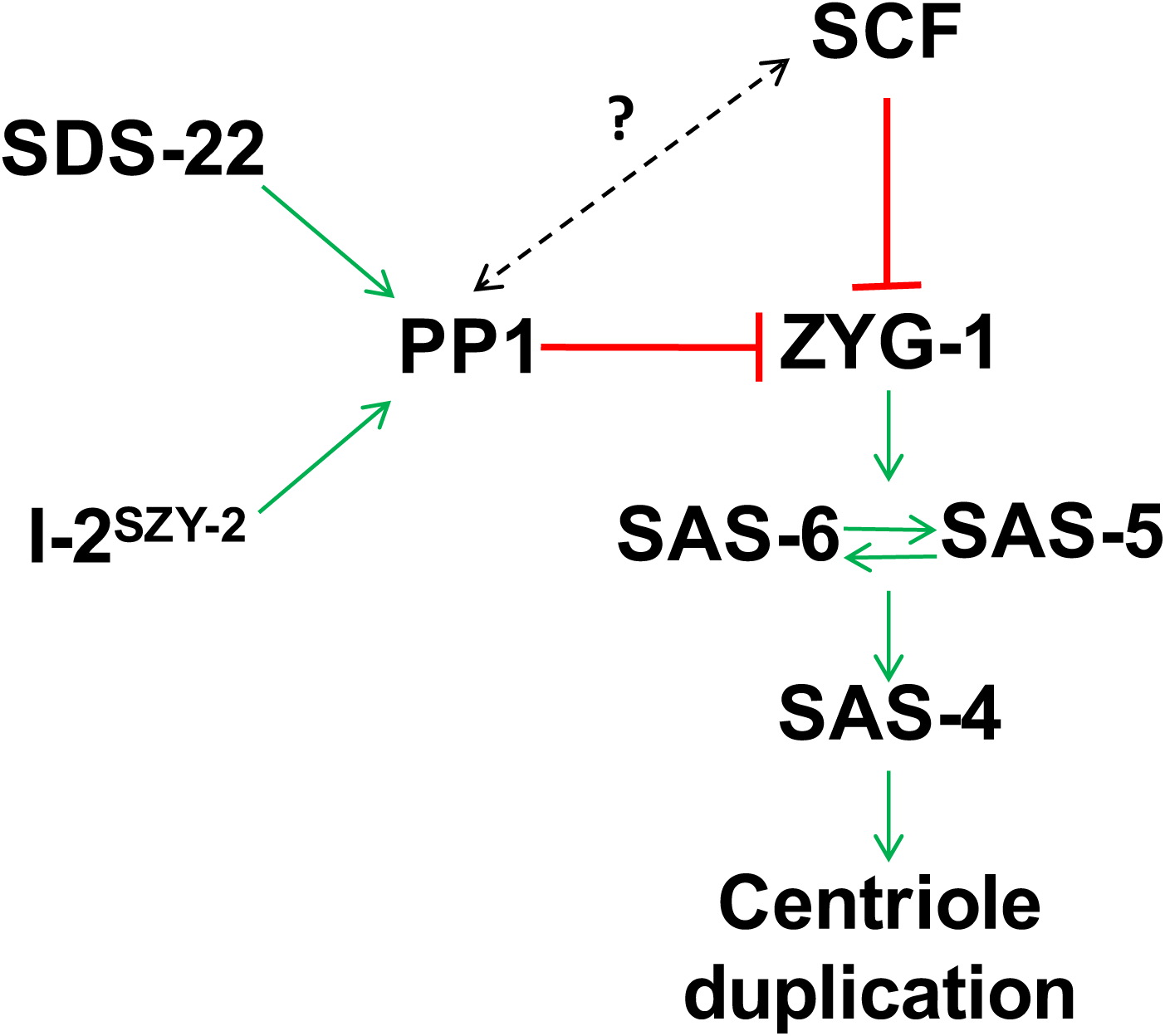
Model of how I-2^SZY-2^, SDS-22 and PP1 might cooperate with the SCF to regulate centriole duplication. Our work shows that PP1, I-2^SZY-2^ and SDS-22 down regulate ZYG-1 levels to constrain centriole duplication. We speculate that this is a new mechanism of regulation that operates independently of the previously documented SCF-mediated regulation. We cannot rule out, however, an interplay between the two pathways (dotted line).

How does PP1 regulate ZYG-1 levels? Although many PP1 targets have been identified none are known regulators of centrosome duplication. A major substrate of PP1, I-2 and SDS22 is Aurora B kinase and activity of the *C. elegans* homolog, AIR-2, is inhibited by PP1 [25,39,43]. Down regulation of AIR-2 activity in the *zyg-1(it25); szy-2(bs4)* double mutant however did not have any measurable effect on the level of centriole duplication suggesting that PP1 is not regulating centriole duplication by antagonizing Aurora B activity. Identical results were obtained from two additional mitotic kinases, Plk1 or Aurora A. Our data indicate that PP1 regulates ZYG-1 levels through a post-translational mechanism. In theory PP1 could regulate the recognition of ZYG-1 by the SCF, however we think that this is unlikely as our results do not match with the canonical mechanism for SCF regulation: substrate recognition by the SCF is regulated by phosphorylation, but the increased phosphorylation associated with PP1 down-regulation would be expected to increase proteosomal degradation, *decreasing* substrate accumulation. We, however, find an *increase* in ZYG-1 levels when PP1 activity is reduced. Furthermore, evidence in *Drosophila* and *C. elegans* indicates that degradation of Plk4/ZYG-1 is antagonized by PP2A [58,59]. Therefore, our favored hypothesis is that PP1 regulates ZYG-1 protein levels through an SCF-independent mechanism, perhaps by removing a stabilizing phosphorylation, however the identity of the opposing kinase remains unknown. We cannot, however, rule out the possibility that PP1 functions through the SCF pathway in a non-canonical fashion to mediate ZYG-1 degradation. Of note, a KVXF consensus site for PP1 binding [60] does exist in ZYG-1, however it is not conserved even in closely related nematode species. Nevertheless, although this is a common motif by which PP1 interacts with its substrates the presence of this motif is not an absolute requirement for PP1 binding.

Our data suggest that PP1, I-2^SZY-2^ and SDS-22 cooperate to regulate overall cellular levels of ZYG-1. Down regulation of PP1 activity leads to accumulation of excess ZYG-1 and in embryos expressing a wild-type version of ZYG-1, the elevated activity is sufficient to drive centriole over-duplication. This is similar to the situation in humans and *Drosophila* where overexpression of homologs of ZYG-1 causes centriole over-duplication [2,5,6]. In agreement with this previous work [2] the appearance of supernumerary centrosomes in *C. elegans* results from the formation of extra daughter centrioles in association with a single mother. Intriguingly, we find that when PP1 activity is decreased, the initial duplication event (that occurs during the first cell cycle following the female meiotic division) is unaffected, yielding bipolar spindles at the two-cell stage. However, beginning with the second centriole duplication event (that occurs at the two cell stage), centrioles commence over-duplicating resulting in multipolar spindles, abnormal divisions, and ultimately lethality.

Why is the first duplication event unaffected by the loss of PP1-mediated regulation while the ensuing duplication events go awry? One unique feature of the first duplication event is that it involves paternally-inherited (sperm) centrioles while later events involve centrioles assembled in the embryo. The sperm-derived centrioles are not, however, permanently immune to elevated ZYG-1 levels, as we have observed cases where, during the second centriole cycle, all four mother centrioles (the two original sperm-derived centrioles plus the two centrioles produced during the first duplication event) produce multiple daughters. Thus, it seems that the elevated level of ZYG-1 present in the embryo only triggers over-duplication of the sperm-derived centrioles following a one-cell-cycle delay. There are at least three possibilities to explain the pattern of over-duplication in PP1-compromised embryos. First, loss of PP1 activity might only lead to overexpression of ZYG-1 beginning with the second centriole duplication event. This however seems unlikely, as we have shown that loss of PP1 activity results in elevated ZYG-1 levels even in the first cell cycle (Fig 4C&D), and that it can suppress the failure of the first duplication event in *zyg-1(it25)* embryos (Fig 1C&D, 2B, 3G). Another possibility is that the critical period for exposure to elevated ZYG-1 is one cell cycle prior to the actual overduplication event. Thus the sperm centrioles, which presumably first encounter elevated ZYG-1 in the zygote, would only over-duplicate during the second centriole cycle. This model is however inconsistent with previous work showing that when *zyg-1* is overactive in the male germ line, centrioles will over-duplicate during spermatogenesis, but when these centrioles are introduced into a wild-type eggs they duplicate normally [61]. Thus the prior exposure of sperm centrioles to elevated ZYG-1 activity does not program them to over-duplicate in the zygote. Finally, a third and more likely possibility is that the centrioles need to be exposed to elevated ZYG-1 for two consecutive cell cycles before they will over-duplicate. Thus sperm-derived centrioles are first exposed to elevated ZYG-1 in the zygote, but do not overduplicate until the second cell cycle, after they have been exposed to elevated ZYG-1 activity for two consecutive cell cycles. There is precedent for this type of regulation; during centriole maturation in human cells, Plk1 activity is needed over the course of two successive cell cycles in order for centrioles to fully mature and become capable of organizing PCM and serving as basal bodies [62]. Thus, this mode of regulation might be a common feature of how polo like kinases operate, at least when it comes to the control of centriole function.

Why do increased ZYG-1 levels lead to centriole overduplication in the *C. elegans* embryo? Normally, levels of centriole-associated Plk4 peak in mitosis before being reduced to a single focus marking the site of procentriole assembly at the G1/S transition [8,20,21]. Centrosome-associated ZYG-1 levels also peak in mitosis [63] and we speculate that elevated ZYG-1 levels prevent it from becoming restricted to a single focus during the ensuing cell cycle, leading to the assembly of multiple daughter centrioles. Ensuring that only a single ZYG-1/Plk4 focus is maintained seems to be a key regulatory step in centriole duplication, however our understanding of how this is achieved remains limited. Clearly excess ZYG-1/Plk4 is sufficient to subvert the normal controls. Since Plk4/ZYG-1 play an important role in the recruitment of SAS-5/ana2/STIL and SAS-6 at the initiation of procentriole assembly [13,14] we envision elevated ZYG-1 levels may be required at this time. However whether continual exposure to elevated ZYG-1 through two cell cycles is required for overduplication remains an open question.

In summary, we have identified I-2^SZY-2^, SDS-22, and PP1 as novel regulators of centriole duplication. The involvement of PP1 in regulating centriole duplication has not previously been described, but interestingly centrosome amplification was reported after depletion of I-2 from human cells, suggesting that the function of PP1 in regulating centriole duplication may indeed be conserved [31]. We show that the key function of PP1 is in limiting the availability of ZYG-1. When PP1 activity is decreased, excessive accumulation of ZYG-1 leads to the formation of extra daughter centrioles in association with a single mother, resulting in centriole amplification. Appropriate regulation of PP1 activity is therefore crucial to maintaining correct centrosome numbers. Although the requirement for PP1 in chromosome segregation due to its function at the kinetochores is well documented, our work suggests a novel requirement for PP1 in the maintenance of genome stability by regulating centriole duplication.

## Experimental Procedures

### Worm Strains and RNAi

All worm strains were maintained at 20°C on MYOB plates seeded with OP50. The strains used in these experiments are listed in supplemental table 1. To monitor the effect of the *sds-22(bs9)* mutation on centriole duplication *sds-22(bs9)* homozygotes were selected from the OC626 strain using the visible *dpy* marker which is closely linked to the *sds-22* gene. RNAi of SZY-2 was carried out by soaking worms in dsRNA [64], all other RNAi experiments were carried out by feeding worms bacteria expressing dsRNA as previously described [65]. Briefly, bacteria containing the RNAi construct were grown overnight and seeded onto MYOB plates supplemented with 25ug/ml ampicillin and 1mM IPTG. For double RNAi a 50:50 mix of overnight cultures of the two bacterial strains was plated. Worms were placed on RNAi at the L4 stage for 28h before analysis. The sequence contained in the GSP-1 RNAi construct was evaluated using the Clone mapper tool (http://bioinformatics.lif.univ-mrs.fr/RNAiMap/index.html)[66], which confirmed likely specificity for only the *gsp-1* gene. For control RNAi we used an *smd-1*-containing vector.

### Immunoprecipitation and Western Blotting

Embryonic extracts for western blots were prepared and analyzed as detailed previously [59]. Whole worm extracts for the immunoprecipitation (IP) experiments were made according to [67]. Total protein concentration was determined using the Biorad Protein Assay Dye Reagent (Bio-Rad). 1.6 mg of total protein from N2 worms was used for performing each IP. Briefly, for each IP, 30 μl of Dynabeads Protein A (Life Technologies) were incubated with 10 μg of each respective antibody at 4°C for 2 hours. Beads were washed three times in worm lysis buffer (50 mM HEPES (pH 7.4), 1 mM EGTA, 1 mM MgCl_2_, 100 mM KCl, 10% Glycerol, 0.05% NP-40), re-suspended in 2X Laemmli Buffer (Bio-Rad), boiled at 100°C for 2 minutes and analyzed by western blotting. The following antibodies/reagents were used in this study: polyclonal SZY-2 antibodies were raised and purified against the entire *szy-2* ORF (Covance); GSP-1 antibodies were raised and purified against the peptide CQYQGMNSGRPAVGGGRPGTTAGKK (YenZym Antibodies LLC); ZYG-1 (Song et. al 2008), phospho-histone H3 (Abcam); GSP-2 [68]; DM1A (Sigma). Mass spectrometry analysis was carried out by the NIDDK mass spec core facility.

### qRT-PCR

RNA was extracted from wild-type and *szy-2(bs4)* embryos, DNase treated and cDNA made using a superscriptIII first strand synthesis kit (Invitrogen). Forward (ACAGTACGCGGAAGAAATGG) and reverse (CACAGCAACCATCTTTTGGA) primers were used to amplify *zyg-1.* Primers against *ama-1* were used as a control [69]. qRT reactions used iQ SYBR green supermix (Bio-Rad) as directed by the manufacturer.

### Imaging

Fixation and staining of embryos was carried out as described previously [36]. The following antibodies were used at a 1/1000 dilution: DM1A (Sigma), phospho-histone H3 (Abcam), ZYG-1 [19], SPD-2 [50] and anti-SAS-4 (Song et. al 2008). For live and fixed imaging we used a spinning disk confocal microscope which has been described previously [61]. To determine whether PP1 affects translation of the ZYG-1 transcript we shifted worms carrying the reporter construct to 25°C as L4s and imaged embryos the next day. Intensities of chromatin GFP at first metaphase were measured using Metamorph. Levels of ZYG-1 or SPD-2 at the centrosome were determined by quantification of average pixel intensity at the centrosome. Maximal projections of the centrosome were used for quantification of fluorescence in ImageJ 1.40g and background fluorescence was subtracted. Centrosome fluorescence was normalized to controls such that control intensity is 1. For structured illumination microscopy, embryos were immuno-labeled as usual and mounted in Vectashield (Vector Laboratories, Inc.). Samples were imaged with a DeltaVision OMX4 SIM Imaging System (Applied Precision).

## Acknowledgements

Some strains were provided by the CGC, which is funded by NIH Office of Research Infrastructure Programs (P40 OD010440). Deletion strains were provided by the National BioResource Project (Japan). We thank M. Colaiacovo for *C. elegans* GSP-2 antibody, and E. Anderson for assistance with mass spectrometry.

## Abbreviations

PP1: protein phosphatase 1

## Supporting Information

**Supplemental Figure S1. Centriole behavior in various genetic backgrounds.** Schematic showing how centrioles behave and how their numbers are established at each cell stage through the first three cell cycles in A) wild-type embryos, B) *zyg-1(it25)* mutant embryos and in C) embryos with reduced PP1 function.

**Supplemental Figure S2. RNAi of PP1 regulators in *zyg-1(it25)* worms.** Quantification of embryonic viability among the progeny of worms grown at the semi-permissive temperature of 24°C and depleted for the indicated PP1 regulators. *smd-1(RNAi)* targets a nonessential gene and serves as a negative control. Only reduction of *sds-22* led to substantial rescue of *zyg-1(it25)* lethality.

**Supplemental Figure S3. Localization of I-2^SZY-2^, PP1β^GSP-1^ and SDS-22.** GFP fusions of the indicated protein were expressed in the embryo. I-2^SZY-2^ Is enriched in the nuclei. Pronuclei meeting during the first cell cycle is shown. PP1β^GSP-1^ is enriched on the chromatin throughout mitosis. Anaphase of the first cell cycle is shown. SDS-22 is weakly localized to the spindle. Metaphase of the first cell cycle is shown.

**Supplemental Figure S4. Summary of Immunoprecipitation-Mass Spectrometry Results.** Each of the indicated proteins was immunoprecipitated from worm extracts and co-purifying proteins identified by mass spec. Shown are the top five hits based on peptide number. In cases where PP1^GSP-1^, SDS-22 or I-2^szy-2^ were not among the top hits, they are also shown along with their rank and number of identifying peptides.

**Supplemental Figure S5. Loss of SDS-22 does not elevate phospho-histone H3 levels in mitotic chromatin.** A) Western blot showing elevated total phospho-histone levels present in extract from mixed stage *sds-22(b9/tm5187)* embryos. This elevation is likely due to an increase in the length of mitosis in *sds-22* mutant embryos as B) quantitative immunofluorescence microscopy shows that mitotic chromatin in *sds-22(bs9/tm5187)* embryos is not enriched for phospho-histone H3 relative to the wild type, and in fact, appears reduced. (a.u. = arbitrary units)

**Supplemental Figure S6. Aurora A^air-1^, Aurora B^AIR-2^, and Plk-1 do not oppose PP1 in the control of centriole duplication.** Each of the specified kinases were RNAi depleted in *zyg-1(i25); szy-2(bs4)* worms and centriole duplication monitored by microscopy. A) Quantification of centriole duplication. Numbers above bars indicate the percentage of successful centriole duplication events and the number within the bars indicate the number of events scored. B-D) Representative stills from timelapse recordings of *zyg-1(it25); szy-2(bs4)* embryos expressing GFP::tubulin and mCherry::histone and treated with control RNAi or RNAi against one of the three indicated mitotic kinases. B) Embryos treated with control RNAi duplicate centrioles, proceed to the 2 cell stage, and build bipolar spindles. C) Embryos treated with *air-2* RNAi duplicate centrioles, but fail in chromosome segregation, thus display 4 centrosomes associated with a single enlarged nucleus. D) Embryos treated with *plk-1* RNAi show a variety of cell division defects yet present with 4 centrosomes indicating centriole duplication has occurred. E&F) Representative stills from time-lapse recordings of *zyg-1(it25); szy-2(bs4)* embryos expressing GFP::SPD-2 and treated with control RNAi (E) or RNAi against *air-1* (F). Centrioles from F are enlarged in a&b. Embryos are at the early 2-cell stage and duplicated centrioles are visible (arrow heads). Note, in this example only one centriole of the control embryo has duplicated.

**Supplemental Figure S7. Centrosome-associated SPD-2 levels are elevated in *szy-2(bs4)* mutants, but this does not appear to contribute to rescue of the *zyg-1(it25)* phenotype.** A) Average GFP::SPD-2 levels at the centrosome during the first cell cycle were calculated and normalized to control. B) Average levels of endogenous SPD-2 at the centrosome during the first metaphase in *szy-2(bs4)* embryos were normalized to controls. C&D) Representative wild type and *szy-2(bs4)* embryos stained for DNA (blue), tubulin (red) and SPD-2 (green). SPD-2 staining at centrosomes is enlarged in a and b. E&F) Comparison of SPD-2::GFP levels at the centrosome in an embryo expressing a codon-optimized version of SPD-2::GFP (OE=overexpression) (E) and a strain expressing GFP::SPD-2 from the native sequence (F). G) Quantification of embryonic viability among the indicated strains to determine whether overexpression of SPD-2 is sufficient to rescue the *zyg-1(it25)* phenotype. In each case n=15 worms, >1000 embryos. H) GFP::SPD-2 levels at the centrosome in *zyg-1(it25); szy-2(bs4)* embryos treated with negative control *smd-1(RNAi)* or *air-1(RNAi).*

**Supplemental Figure S8. SAS-6 levels are not altered by reduction of PP1 activity.** A) Western blot demonstrating equivalent total levels of SAS-6 in wild-type and *szy-2(bs4)* embryos. B) Quantitation of western blot data (n=2). C) Average levels of centrosome-localized GFP::SAS-6 in wild-type and *szy-2(bs4)* embryos were determined throughout the first cell cycle. Values shown are normalized to the average wild-type value at each stage. Error bars represent standard error.

**Supplemental Figure S9. Centrioles continue to over-duplicate in older embryos.** Select frames from a time-lapse recording of an *sds-22(bs9/tm5187)* embryo expressing GFP::SPD-2 and mCherry::histone. The arrowhead shows a spindle pole (frame 18:00) giving rise to multiple centrosomes during the next cell cycle (fram 28:00). White arrows indicate centrosomes that become inactive during the round of division following their birth (frames 10:00 and 18:00) but become active again during the next cell cycle (frames 22:00-34:00). Note that both centrosomes were not always visible in these frames.

### Supplementary Movies

**Movie S1-** Normal centriole duplication in a wild-type embryo expressing GFP::tubulin and mCherry::histone. Note bipolar spindle formation at the two-cell stage.

**Movie S2-** Centriole duplication in a *zyg-1(it25)* mutant embryo expressing GFP::SPD-2 and mCherry::histone. Note monopolar spindle formation at the two-cell stage.

**Movie S3-** Centriole duplication in *zyg-1(it25); szy-2(bs4)* double mutant embryo expressing GFP::tubulin and mCherry::histone. Note restoration of bipolar spindle formation at the two-cell stage.

**Movie S4-** Anaphase bridges can be seen in embryos expressing GFP::histone and mCherry::SPD-2 when *gsp-1 and gsp-2* are co-depleted by RNAi.

**Movie S5-** Embryos expressing GFP::histone and mCherry::SPD-2 produce extra centrosomes at the 4cell stage when *gsp-1 and gsp-2* are co-depleted by RNAi.

**Movie S6-** Extra centrosomes are present at the four-cell stage in *sds-22(bs9/tm5187)* embryos expressing GFP::SPD-2 and mCherry::histone.

**Movie S7-** In *sds-22(bs9/tm5187)* embryos expressing GFP::SPD-2 and mCherry::histone, extra centrosomes continue to be produced after the 4-cell stage. The extra centrosomes recruit PCM and participate in spindle assembly.

**Movie S8-** The extra centrosomes detected in *gsp-1(RNAi); gsp-2(RNAi)* embryos expressing GFP::histone and mCherry::SPD-2 function as microtubule-organizing centers and give rise to multipolar spindles.

**Supplemental Table 1.** C.elegans strains used in this study

| Strain name | Genotype |
| --- | --- |
| OC467 | *szy-2(tm3972)/hT2[bli-4(e937) let?(q782) qIs48](I;III)* |
| OC192 | *szy-2(bs4) III* |
| OC227 | *zyg-1(it25) II; szy-2(bs4) III* |
| OC14 | *zyg-1(it25) II* |
| OC192 | *szy-2(bs4) III* |
| OC199 | *zyg-1(it25) sds-22(bs9) II* |
| OC367 | *zyg-1(it25) II; szy-2(bs4) unc-119(ed3) III; ltIs37[pAA64: unc-119(+) pie-1p::mcherry::his58]; ltIs25[pAZ132: unc-119(+) pie-1p::gfp::tba-2]* |
| OC429 | *dpy-17(e164) szy-2(bs4) III; bsls2 [pCK5.5: : unc-119(+) pie-1p::gfp::spd-2]* |
| OC448 | *zyg-1(it25) II; ltIs37[pAA64: unc-119(+) pie-1p::mcherry::his-58]; ltIs25[pAZ132: unc-119(+) pie-1p::gfp::tba-2]* |
| OC469 | *zyg-1(it25) II; szy-2(tm3972) III; ltIs37[pAA64: unc-119(+) pie-1p::mcherry::his-58]; ltIs25[pAZ132: unc-119(+) pie-1p::gfp::tba-2]* |
| OC573 | *sds-22(tm5187)/mIn1[dpy-10(e128) mIs14(myo-2::gfp)] II* |
| OC156 | *zyg-1(it25) II; bsIs2 [pCK5.5: : unc-119(+) pie-1p::gfp::spd-2]; unc-119(ed3) ruls32[unc-119(+) pie-1p::gfp::his-58] III* |
| OC381 | *zyg-1(it25) II; gsp-2(tm301)/ hT2[bli-4(e937) let?(q782) qIs48](I;III); ItIs37[pAA64: unc-119(+) pie-1p::mcherry::his-58]; ItIs25[pAZ132: unc-119(+) pie-1p::gfp::tba-2]* |
| OC626 | *sds-22(bs9)/sds-22(tm5187) II; bsIs2 [pCK5.5: unc-119(+) pie-1p::gfp::spd-2]; ItIs37 [pAA64: unc-119(+) pie-1p::mcherry::his-58]* |
| OC591 | *zyg-1(it25); [unc-119(+) pie-1-GFP::SPD-2OE]Insertion from Decker et al, 2011* |
| OC92 | *unc-119(ed4) III; bsIs2 [pCK5.5: unc-119(+) pie-1p::gfp::spd-2]* |
| NIN33 | *szy-2(bs4) unc119(ed3) III; fem-1(hc17) IV; ItIs33[pOD224: unc-119(+) pie-1p::gfp::tev::stag::sas-6;]* |
| OC770 | *bsSi15 [pKO109: spd-2p::spd-2::mCherry::spd-2 3'-utr, unc-119(+)] I; szy-2(tm3972) III* |
| OC779 | *bsSi15 [pKO109: spd-2p::spd-2::mCherry::spd-2 3'-utr, unc-119(+)] I; bsSi30[pCW9: ZC504.3p::sfGFP::his-58::ZC504.3 3'-utr, unc-119(+)] II* |
| OC723 | *bsSi9 [pKO113: unc-119(+) zyg-1p::gfp::his-58::3'utr zyg-1] I; unc-119(ed3) III* |
| OC835 | *bsIs9 [pKO113: unc-119(+) zyg-1p-gfp::his-58-zyg-1 3'utr]I; szy-2(tm3972) III* |
| OC547 | *unc-119(ed3)III: bsIs33 [pNP121: pie-1promoter-gfp-sds-22-pie-1UTR unc-119(+)]/+* |
| OC484 | *unc-119(ed3) III; BsIs18[pNP97: pie1promoter-gfp-gsp-1genomic-pie1UTR unc-119(+)]* |
| OC426 | *unc119(ed3)III; pNP71: bsIs52[pie-1promoter-gfp::szy-2-pie-1UTR unc119(+)]* |

